# Ubiquitin Proteasome System Components, RAD23A and USP13, Modulate TDP-43 Solubility and Neuronal Toxicity

**DOI:** 10.1101/2025.02.06.636899

**Authors:** Casey Dalton, Jelena Mojsilovic-Petrovic, Nathaniel Safren, Carley Snoznik, Kamil K. Gebis, Yi-Zhi Wang, Todd Lamitina, Jeffrey N. Savas, Robert G. Kalb

## Abstract

At autopsy, >95% of ALS cases display a redistribution of the essential RNA binding protein TDP-43 from the nucleus into cytoplasmic aggregates. The mislocalization and aggregation of TDP-43 is believed to be a key pathological driver in ALS. Due to its vital role in basic cellular mechanisms, direct depletion of TDP-43 is unlikely to lead to a promising therapy. Therefore, we have explored the utility of identifying modifier genes that modify its mislocalization or aggregation. We have previously shown that loss of *rad-23* improves locomotor deficits in TDP-43 *C. elegans* models of disease and increases the degradation rate of TDP-43 in cellular models. To understand the mechanism through which these protective effects occur, we generated an inducible mutant TDP-43 HEK293 cell line. We find that knockdown of *RAD23A* reduces insoluble TDP-43 levels in this model and primary rat cortical neurons expressing human TDP-43^A315T^. Utilizing a discovery-based proteomics approach, we then explored how loss of *RAD23A* remodels the proteome. Through this proteomic screen, we identified USP13, a deubiquitinase, as a new potent modifier of TDP-43 induced aggregation and cytotoxicity. We find that knockdown of *USP13* reduces the abundance of sarkosyl insoluble mTDP-43 in both our HEK293 model and primary rat neurons, reduces cell death in primary rat motor neurons, and improves locomotor deficits in *C. elegans* ALS models.

**Significance Statement:** Amyotrophic lateral sclerosis (ALS) is a fatal neurodegenerative disease (NDD) with no effective therapies. The mislocalization and aggregation of TAR DNA binding protein 43 (TDP-43) is a key pathological marker of ALS and other NDDs. Due to its vital functions, targeted therapeutic reduction of TDP-43 could be problematic. Here, we have explored the utility of targeting modifier genes. We find that knockdown of two members of the ubiquitin proteasome system, *RAD23A* and *USP13*, enhance TDP-43 solubility and decrease TDP-43 induced neurotoxicity.

## Introduction

Amyotrophic lateral sclerosis (ALS) is a progressive, neurodegenerative disease characterized by the loss of both upper and lower motor neurons resulting in muscle denervation, weakness, ventilatory failure, and death typically within 3 to 5 years of diagnosis (1). Approximately 10-15% of ALS cases have a monogenic etiology (familial ALS or fALS) and the remainder are classified as sporadic (or sALS) (2). fALS and sALS are clinically indistinguishable and share neuropathological features supporting the view that inquiry into fALS pathophysiology will be relevant to all individuals with ALS. A key pathological hallmark of ALS is the accumulation of ubiquitinated, misfolded, detergent-insoluble proteins (3), likely reflecting dysfunction of cellular proteostasis machinery. The central role of the proteostasis network in ALS pathogenesis is further reinforced by the discovery of mutations in genes within the ubiquitin proteasome system (UPS) and lysosome-autophagy pathway (*UBQLN2*, *SQSTM1*, *HSP17*, and others) that can cause fALS.

One protein of particular interest is TDP-43. TDP-43 is aggregated, ubiquitinated, hyperphosphorylated and mislocalized in the cytoplasm in the brain and spinal cord of >95% of ALS patients, ∼45% of FTD patients and up to 74% of Alzheimer’s patients at autopsy (4–8). TDP-43 likely contributes to pathophysiological events by both loss-of-function (i.e., nuclear depletion) and gain-of-function (i.e., cytoplasmic aggregation) mechanisms. Missense mutations in TDP-43 have been identified in a small number of fALS cases (9). These mutations increase aggregation propensity, aggravate cytoplasmic mislocalization, disrupt physiological interactions and influence stability leading to resistance to degradation (10–12). A novel solubilization method from the Polymenidou lab, SarkoSpin, shows that aggregated, phosphorylated, ubiquitinated TDP-43 can be isolated from patient tissue and induce cytotoxicity when applied to naïve neurons (13). This work separates toxic aggregated TDP-43 from physiological TDP-43 and protein complexes after tissue lysis in sarkosyl and benzonase, leaving the cytotoxic aggregated TDP-43 in the sarkosyl insoluble fraction. Thus, sarkosyl insoluble TDP-43 is either a toxic entity or reproducibly co-purifies with (and is a reporter for) the toxic species.

Due to its vital physiological functions, direct depletion of TDP-43 is not a promising therapeutic approach. Therefore, we have explored the utility of identifying and targeting modifier genes that ameliorate TDP-43 toxicity. In a forward genetic screen in two *Caenorhabditis elegans* (*C. elegans)* models of fALS, loss of *rad-23* improved mutant TDP-43-induced and mutant SOD-induced locomotor deficits (14). Knockdown (KD) of *RAD23* in mammalian neurons also reduces SOD1- and TDP-43-induced toxicity and enhanced turnover of these substrates in HEK293 cells. RAD23 was originally identified in yeast during a screen for modifiers of UV irradiation sensitivity (15) and is now recognized as a component of the nucleotide excision repair pathway. In addition, RAD23 functions as a shuttle factor that binds ubiquitinated substrates and presents them to the proteasome for degradation (16, 17). The protein exists as a single isoform in *Saccharomyces cerevisiae* and *C. elegans,* and as two isoforms in vertebrates, RAD23A and RAD23B. In addition to promoting protein degradation, several studies have shown that RAD23 can also stabilize certain substrates such as XPC (18), p53 (19), and TDP-43 (14).

In this study we sought to identify which RAD23 isoform is responsible for the observed protective effects of RAD23 knockdown and the mechanism through which these effects occur. Using tandem mass tag-based quantitative proteomics we show how RAD23 exerts broad control over the composition of the proteome and we identify USP13 as a new potent modifier of TDP-43-induced pathology.

## Results

### Knockdown of *RAD23A* but not *RAD23B* reduces insoluble TDP-43

We generated doxycycline (DOX)-inducible hemagglutinin (HA)-tagged Q331K human TDP-43 (+mTDP-43) expressing cells using the Flp-In T-Rex HEK293 cell line. This system allows temporal control of TDP-43 expression. Following a doxycycline pulse-chase, we find a HA-mTDP-43 half-life of ∼31 hours which is consistent with prior studies using inducible cell lines (20). We next asked whether induced mTDP-43 enters the insoluble fraction of cell lysates and whether KD of *RAD23A*, *RAD23B* or both are required for this observation. To accomplish this, we utilized SarkoSpin fractionation, with siRNA KD of each isoform of RAD23. We find mTDP-43 in the sarkosyl insoluble fraction following 24 hours of +mTDP-43 pulse. In these cells, RNAi mediated *RAD23A* KD achieves an average of ∼61% reduction in RAD23A and ∼23% increase in RAD23B while *RAD23B* KD achieves an average of 84% reduction in RAD23B and 19% reduction in RAD23A (Fig. 1A). Notably, we find that KD of *RAD23A* but not *RAD23B* reduces the percentage of sarkosyl insoluble mTDP-43 by ∼33%, indicating it is KD of the RAD23A isoform that increases TDP-43 solubility (Fig 1B and C). We next asked if KD of *RAD23A*, *RAD23B*, or both could affect the cytoplasmic mislocalization of mTDP-43. We consistently find that ∼30-35% of HA-mTDP-43 exhibits cytoplasmic mislocalization 24 hours after doxycycline induction (SI Appendix, Fig. S1). KD of either *RAD23A* (∼11%) or *RAD23B* (∼19%) significantly reduces the amount of cytosolic mTDP-43; however, neither condition results in a significant reduction in the percentage of cytosolic TDP-43 (SI Appendix, Fig. S1). This result suggests KD of either isoform may produce a modest increase in total TDP-43 degradation; however, KD of neither isoform affects the cytoplasmic mislocalization phenotype. Taken with our solubility studies, we determined it was likely KD of *RAD23A* driving protective effects through reductions in insoluble TDP-43. To determine the effects of complete elimination of RAD23A, we knocked out *RAD23A* using CRISPR Cas9 in our HA-mTDP-43 inducible cell line. Knockout further reduces the percentage of insoluble TDP-43 to ∼50% of the control (Fig. 1D, E, and F). Together these results demonstrate a dose-dependent effect of RAD23A protein on the abundance of sarkosyl insoluble TDP-43.

**Figure 1.**
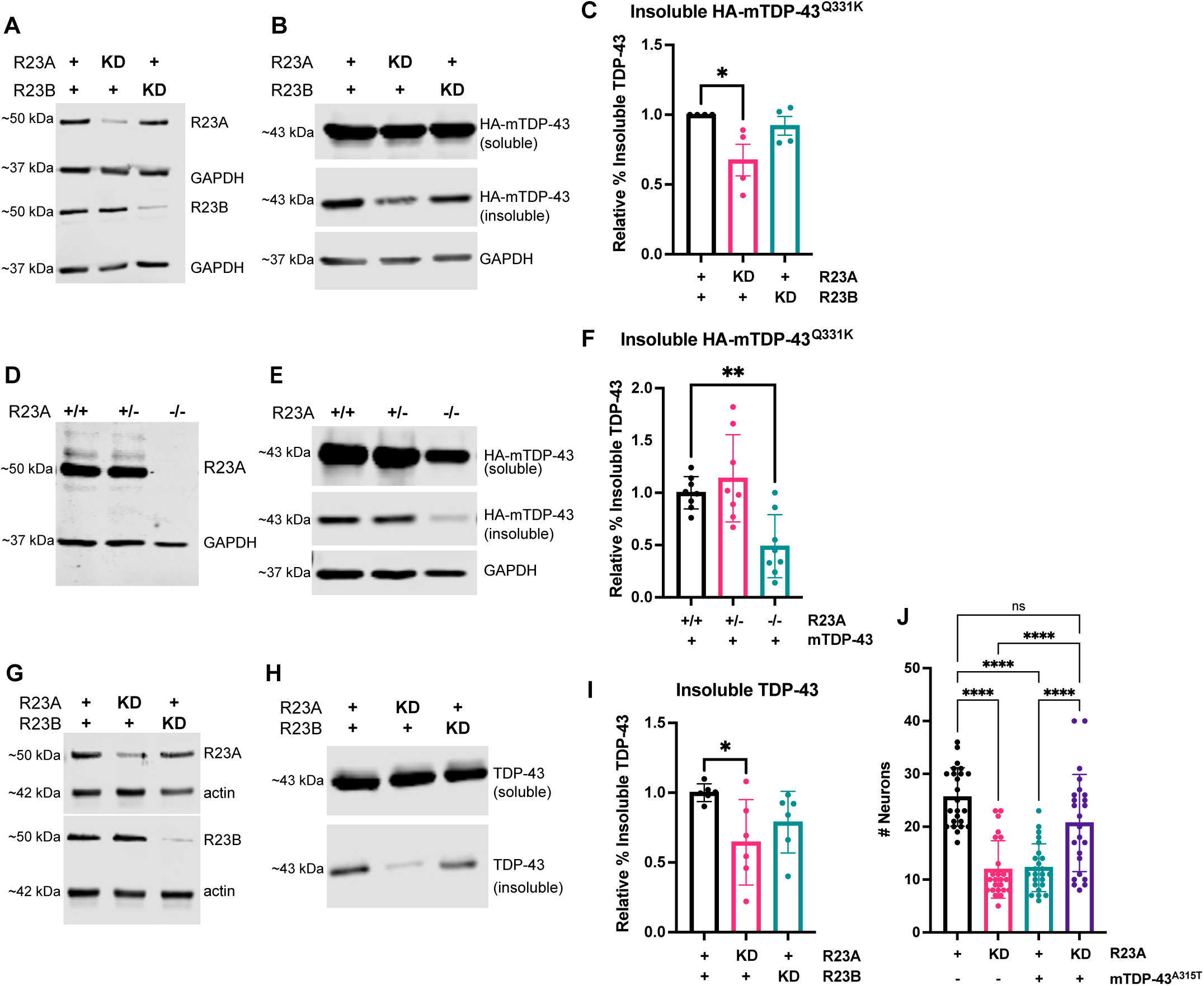
Loss of *RAD23A* reduces insoluble TDP-43 and blunts TDP-43 induced neuronal toxicity. **A - C:** Inducible HA-mTDP-43^Q331K^ Flp-In T-REx HEK293 cells were plated, transfected with indicated siRNA, induced with DOX for 24hrs, lysates collected and subjected to SarkoSpin fractionation. **(A)** *RAD23A* and *RAD23B* siRNA efficiency and isoform specificity, representative immunoblot. **(B)** HA-mTDP-43 soluble and insoluble fractions, representative immunoblot. **(C)** *RAD23A* knockdown reduces the percentage of HA-mTDP-43 present in the insoluble fraction (quantified as a percent of total TDP-43) relative to negative control. Data represent the mean +/−SEM, N=4. (one way ANOVA, F(2,9)=4.97, *p<0.05). **D - F:** R23A^+/+^, R23A^+/−^, R23A^−/−^ CRISPR generated knockout in HA-mTDP-43 Flp-In T-REx HEK293 cells were plated, induced with DOX for 24hrs, lysates collected and subjected to SarkoSpin fractionation. **(D)** RAD23A expression levels, representative immunoblot **(E)** Insoluble and soluble HA-mTDP-43, representative immunoblot. **(F)** R23A^−/−^ knockout shows a reduction in HA-mTDP-43 present in the insoluble fraction (quantified as a percent of total TDP-43) expressed as a fold-change. Data represent the mean +/− SD, N=4, two technical replicates per experimental replicate. (one-way ANOVA, F(2,21)=9.728, **p<0.01) **G - I :** Primary rat cortical neuron cultures were transfected with indicated siRNAs on DIV5, infected with HSV-mTDP-43^A315T^ on DIV12, and collected after 48 hrs for SarkoSpin fractionation. **(G)** *RAD23A* and *RAD23B* siRNA efficiency and isoform specificity, representative immunoblot **(H)** TDP-43 soluble and insoluble fractions, representative immunoblot. **(I)** *RAD23A* knockdown reduces TDP-43 in the insoluble fraction (quantified as a percent of total TDP-43) relative to the negative control. Data represent the mean +/−SEM, N=3, two technical replicates per experimental replicate. (one way ANOVA, F(2,15)=3.931, *p<0.05). **(J)** Primary rat motor neuron survival following HSV-mTDP-43^A315T^ expression and *RAD23A* knockdown. Primary rat cortical neuronal cultures transfected with indicated siRNAs on DIV5, infected with HSV-mTDP-43 or HSV-LacZ on DIV14, and fixed and stained with motor neuron marker SMI-32 on DIV19. N=4 experimental replicates, each data point is an area counted, N=6 areas per replicate. (one way ANOVA, F(5,138)=35.03, ****p<0.0001).

To test the effects of loss of RAD23 on insoluble TDP-43 in a more disease relevant cell type, we conducted SarkoSpin fractionation in primary rat cortical neurons transduced with a recombinant HSV viral vector driving the expression of human mTDP-43^A315T^. In this neuronal system, RNAi KD of *RAD23A* achieved an average of ∼52% reduction in RAD23A and ∼14% reduction in RAD23B, while KD of *RAD23B* produced an average of ∼68*%* reduction in RAD23B with ∼5% reduction in RAD23A (Fig. 1G). KD of *RAD23A*, but not *RAD23B*, reduced the percentage of insoluble TDP-43 by ∼36%. (Fig. 1H and I). Finally, we asked if reducing RAD23A would protect neurons from TDP-43 induced neurotoxicity. To test this, we conducted a viability assay in primary rat motor neurons expressing mTDP-43^A315T^ and found that *RAD23A* KD increases survival in neurons expressing mTDP-43^A315T^ by ∼69% (Fig. 1J). In summary, we find that loss of *RAD23A* significantly reduced insoluble TDP-43 in a dose-dependent manner in our inducible mTDP-43 model. In addition, *RAD23A* KD significantly reduced sarkosyl insoluble TDP-43 and reduced motor neuron death in primary rat neurons expressing mTDP-43.

### TDP-43 induced proteome remodeling is partially blunted by RAD23A knockdown

RAD23 can act to both increase stability of some substrates and facilitate the degradation of other substrates (14, 16–18). We hypothesized that KD of *RAD23A* would lead to increased expression of proteins that enhance TDP-43 solubility and/or degradation and downregulation of proteins that reduce solubility. To test this hypothesis, we utilized an unbiased proteomics approach by conducting tandem mass tag (TMT)-based quantitative proteomic analysis of our HA-mTDP-43 expressing cells. A total of four experimental groups were analyzed: 1) no mTDP-43 induction without *RAD23A* KD (no mTDP-43;R23A^cont^), 2) no mTDP-43 induction with *RAD23A* KD (no mTDP-43; R23A^KD^), 3) +mTDP-43 induction without *RAD23A* KD (+mTDP-43; R23A^cont^) and 4) +mTDP-43 with *RAD23A* KD (+mTDP-43; R23A^KD^). We focused on three comparison groups: 1) proteins altered by the induction of mTDP-43 (no mTDP-43; R23A^cont^ vs +mTDP-43; R23A^cont^); 2) proteins altered by *RAD23A* KD (no mTDP-43; R23A^cont^ vs no mTDP-43; R23A^KD^) and 3) proteins altered by *RAD23A* KD in the presence of mTDP-43 (+mTDP-43; R23A^cont^ vs +mTDP-43; R23A^KD^).

In total over 6000 proteins were quantified in at least two replicates in each condition. After calculating the log_2_FC and conducting a one-tailed t-test, values were screened with Benjamini-Hochberg (BH) correction. Although no proteins passed BH correction, we find significant enrichment when conducting pathway analysis. We moved forward using the data set as a discovery platform to identify potential proteins and pathways of interest. For initial group comparisons and pathway analysis, proteins with a log_2_FC > 0.25 or < −0.25 and a p-value <0.05 were included. Group 1 contains 281 proteins with significant fold change, of which 171 proteins (61%) have increased expression and 110 (39%) have decreased expression (Fig. 2A and C)(SI Appendix, Table S1), Group 2 contains 219 proteins with significant fold change, of which 139 (63%) have increased expression and 80 (37%) have decreased expression (Fig. 2A and D). Group 3 contains 337 proteins with significant fold change. In contrast to the other two groups, the majority of proteins in Group 3 have decreased expression (74%) while only 87 proteins (26%) have increased expression (Fig. 2A and E). When analyzing these 337 proteins identified in Group 3, against a control, we find that generally *RAD23A* KD opposes the alterations in protein expression evoked by mTDP-43 induction (Fig. 2B).

**Figure 2.**
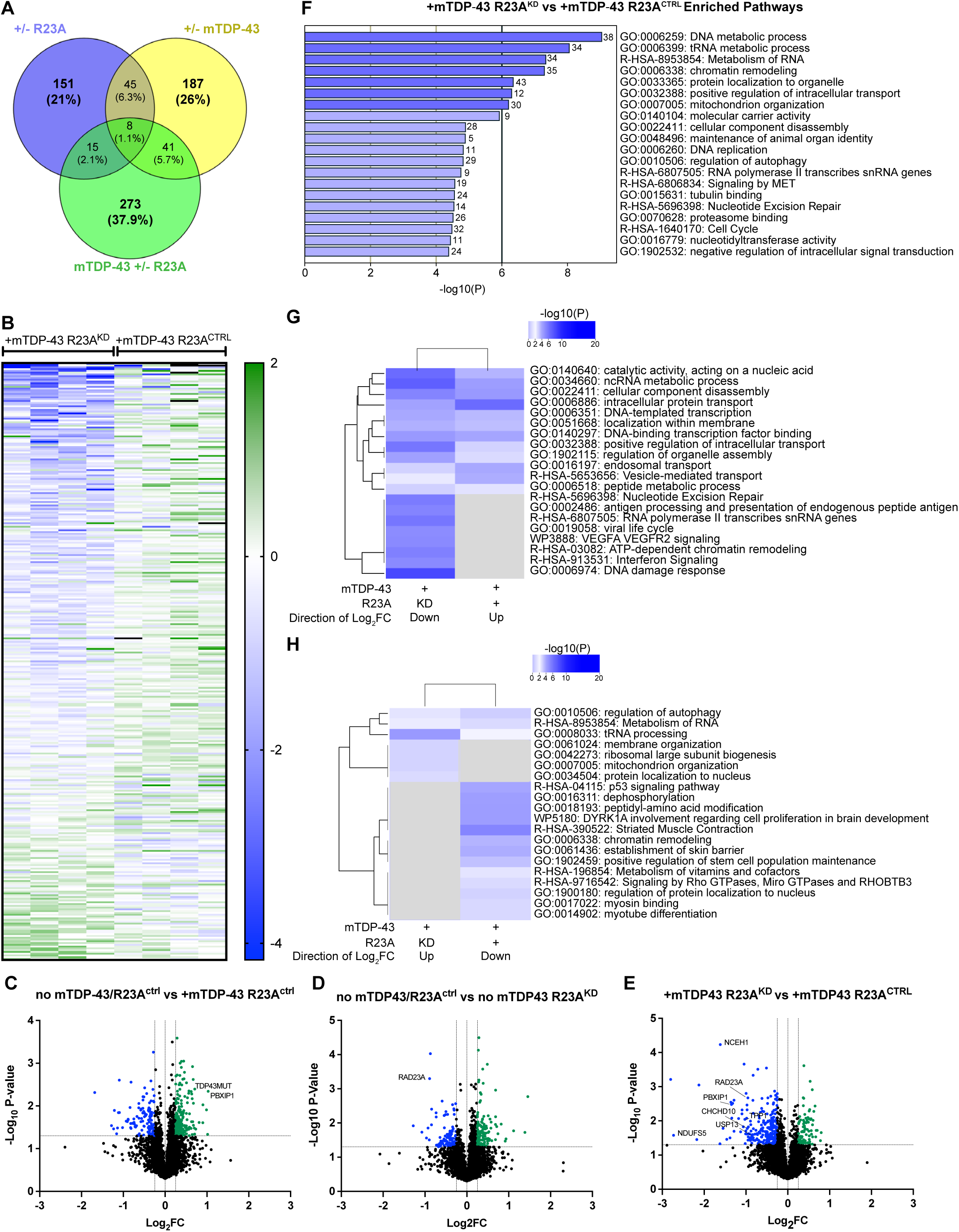
Effects of *RAD23A* knockdown on mTDP-43-induced proteomic remodeling. **(A)** Venn diagram showing the shared proteins between the three comparison groups: Group 1, proteins altered by the induction of mTDP-43, Group 2, proteins altered by *RAD23A* knockdown and Group 3, proteins altered by *RAD23A* knockdown in the presence of mTDP-43. **(B)** Heat Map showing the relative Log_2_FC expression of Group 3 proteins (altered by *RAD23A* knockdown in the presence of mTDP-43) +mTDP-43; R23A^cont^ vs +mTDP-43; R23A^KD^ expressed versus a negative control (no mTDP-43; R23A^cont^). Group 3 proteins shown are those with P<0.05,one-tailed t-test, N=4 replicates, Log_2_FC>0.25 or <-0.25. **C – E.** Volcano plots of protein expression changes in each comparison group. Highlighted data points are those with P<0.05,one-tailed t-test, N=4 replicates, Log2FC>0.25 or <-0.25. **(C)** Volcano Plot of Group 1 proteins, altered expression by mTDP-43 induction. (**D)** Volcano Plot of Group 2 proteins, altered expression by *RAD23A* knockdown. (**E)** Volcano Plot of Group 3 proteins, altered expression by *RAD23A* knockdown in the presence of mTDP-43. (**F)** Pathways enriched in Group 3 using Metascape gene analysis, both up and down regulated proteins were analyzed together. **G, H.** Comparison of shared enriched pathways between Group 1 and Group 3. (**G)** The pathways of upregulated proteins in Group 1 compared to the pathways down regulated proteins in Group 3. (**H)** The pathways of downregulated proteins in Group 1 compared to the pathways of upregulated proteins in Group 3.

The shared proteins between groups yield further insight. 49 proteins are common to both Group 1 (altered by mTDP-43 induction) and Group 3 (altered by *RAD23A* KD in the presence of mTDP-43). 35 proteins that display increased expression by +mTDP-43 induction are decreased following *RAD23A* KD in the presence of mTDP-43, and 12 proteins with decreased expression following +mTDP-43 induction are increased following *RAD23A* KD in the presence of mTDP-43 induction. Only 2 are not altered in opposing directions (SI Appendix, Fig. S2). Surprisingly, Groups 1 and 2 share more common proteins than any other pair at 53. Interestingly, all of the common proteins have differential expression in the same direction (SI Appendix, Fig. S2). This suggests the effects of *RAD23A* KD on the proteome fundamentally depend on whether mTDP-43 is co-expressed. This is further highlighted when comparing Group 2 and Group 3 where there are 23 common proteins, of which 17 move in opposing directions while 6 are altered in the same direction (SI Appendix, Fig. S2). Overall, these results highlight the context dependent effects RAD23A has on the proteome and support the hypothesis that loss of *RAD23A* reverses some of the proteomic alterations evoked by mTDP-43 expression.

Next, we utilized Metascape gene annotation and analysis resources (21), to determine if our differentially expressed proteins were enriched in any specific pathways (SI Appendix, Table S2). For Group 1, pathways typical of TDP-43 pathology such as autophagy, RNA metabolism, chromatin remodeling, and protein localization were enriched in our analysis (20, 22–25) (SI Appendix, Fig. S3). In Group 3, pathways related to RAD23A function such as the DNA damage pathway (18) and proteasome binding (16) as well as some known TDP-43 pathways such as autophagy, RNA metabolism, and protein localization are enriched (Fig. 2F). Group 2 is similarly enriched in pathways related to canonical RAD23A function such as the DNA-repair metabolic process, regulation of protein stability, and deubiquitination; however, there are clear divergences from Group 3 with enrichment in pathways such as apoptosis modulation (SI Appendix, Fig. S3).

When analyzing shared pathways between groups, we find that Groups 1 and 2 share many common pathways, both when analyzing up and down regulated proteins together and when looking at each set separately (SI Appendix, Fig. S4). In contrast, when we compare Group 3 to Groups 1 and 2, while there are shared pathways, if up and down regulated proteins are analyzed separately very few commonalities remain (SI Appendix, Fig. S4). For Groups 1 and 3, similar to the protein level, pathways such as intracellular protein transport and DNA binding transcription factor binding that are upregulated after mTDP-43 induction are downregulated by *RAD23A* KD in the presence of mTDP-43, and to a lesser extent pathways such as autophagy and metabolism of RNA that are downregulated by TDP-43 induction are upregulated by *RAD23A* KD (Fig. 2G and H). Overall, these results are consistent with our hypothesis that *RAD23A* KD blunts TDP-43 induced proteome remodeling, at both the protein and pathway level.

### Protein Candidate Screening

Proteins that change expression in a RAD23A dependent manner may mediate the effects of RAD23A on TDP-43 toxicity. We created a list of candidate modifier genes from Group 3 (+mTDP-43; R23A^cont^ vs +mTDP-43; R23A^KD^) of our mass spectrometry analysis based on: 1) proteins that were identified in our pathway analysis, 2) total fold change, 3) presence in the nervous system based on the Human protein atlas (26), and 4) association with neurodegenerative disease (Table 1). Because most of our top hits were downregulated following mTDP-43 induction and *RAD23A* KD, our best candidates were in this subset of proteins. In an initial screen of these candidates, we conducted a Live-Dead viability/cytotoxicity plate reader assay^®^ in our inducible mTDP-43 cells with saturated doxycycline induction. From this screen, we determined siRNA KD of two candidates (*PBXIP1* and *CHCHD10*) decreased cell viability, while the remaining candidates had no significant impact on cell viability. We then utilized SarkoSpin fractionation following siRNA KD of candidates (SI Appendix, Fig. S5). siRNA KD of three candidates (*USP13*, *NCEH1*, *NDUFS5*) significantly reduced the percent of insoluble TDP-43 (SI Appendix, Fig. S5). Ubiquitin specific peptidase 13 (USP13) is a deubiquitinase implicated in cancer prognosis, neurodegenerative disease and regulation of critical pathways such as autophagy, ER-associated degradation, and the DNA-damage response (27–29). Neutral cholesterol esterase hydrolase 1 (NCEH1) is a hydrolase involved in lipid metabolism (30). NADH Dehydrogenase (Ubiquinone) Iron-Sulfur Protein 5 (NDUFS5) is subunit of complex 1 in the electron transport chain (31). Due to its connection to the protein homeostasis machinery, we decided to further investigate the effect of *USP13* KD on TDP-43 aggregation and toxicity.

**Table 1:**
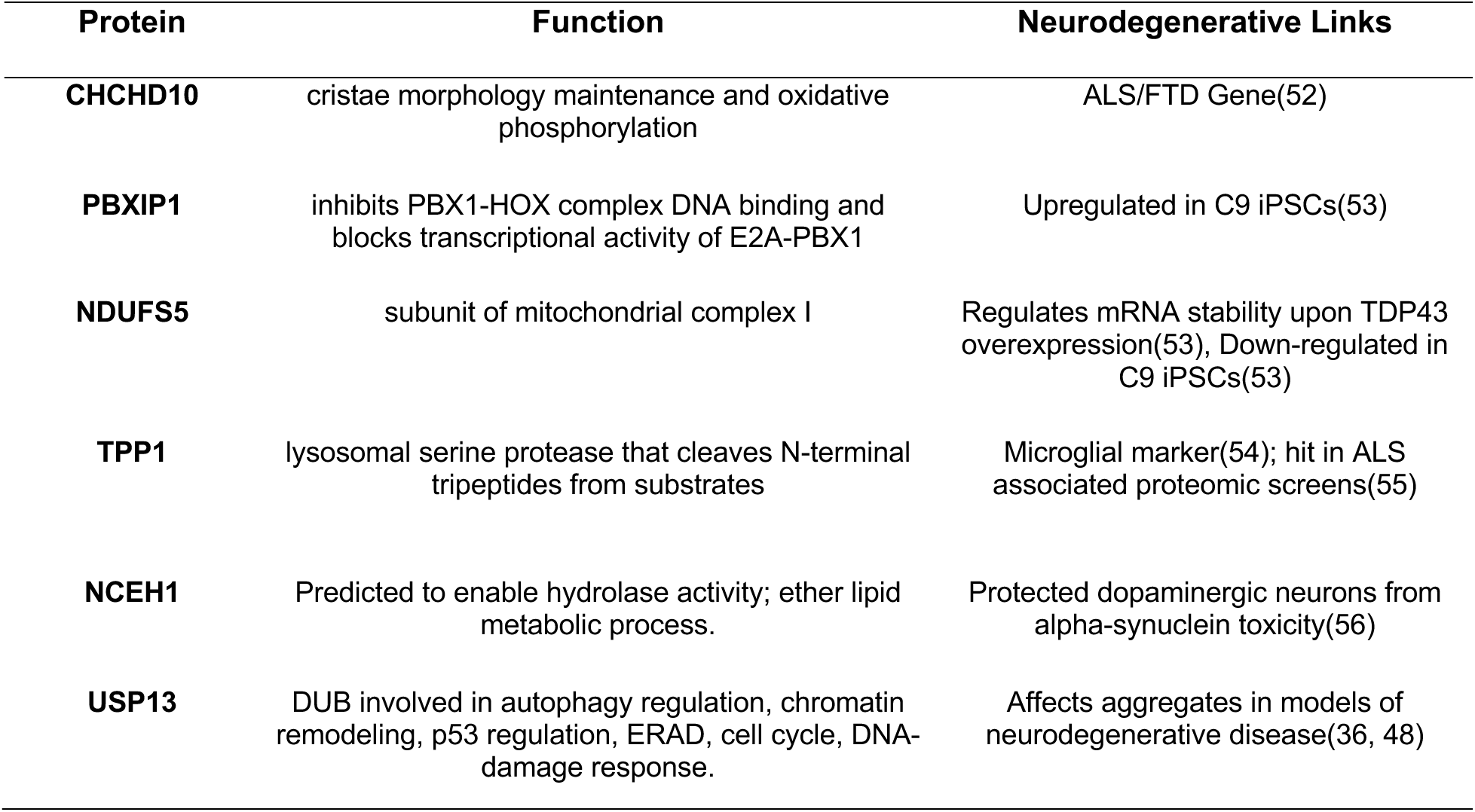
Proteomic candidates function and neurodegenerative links from literature review.

### USP13 is modulated by RAD23A and effects TDP-43 solubility through a distinct pathway

Our TMT proteomics data (Fig. 3A) indicates that *RAD23A* KD in the presence of mTDP-43 induction reduces the amount of USP13. Here we confirm this result by western blot where we see that USP13 is significantly increased by mTDP-43 induction and decreased by mTDP-43 induction with *RAD23A* KD (Fig. 3B and C). Because *RAD23A* KD leads to a reduction in USP13, we hypothesized that USP13 abundance may be regulated by the presence of a USP13/RAD23A complex. To test this, we heterologously expressed HA-RAD23A and conducted immunoprecipitation (IP) of HA-RAD23A and immunoblotted for USP13. We immunoprecipitate HA-RAD23A but not USP13, indicating that the effect of *RAD23A* KD on the abundance of USP13 is likely not due to a direct physical interaction (Fig. 3D). To gain further insight into how RAD23A and USP13 may interact, we asked if their actions on TDP-43 were additive. To do this we first confirmed the results of our candidate screen and find that *USP13* KD reduces the percentage of insoluble TDP-43 in our inducible HA-mTDP-43 cell line by ∼42% (Fig. 3E and F). We next asked if *USP13* KD could further reduce insoluble TDP-43 in our R23A^−/−^ mTDP-43 model. We find that *USP13* KD reduces insoluble TDP-43 by an additional 21% (Fig. 3G and H). These results suggest that while *RAD23A* KD can affect USP13 abundance, they potentially modulate TDP-43 solubility through converging parallel mechanisms rather than through a sequential shared pathway.

**Figure 3.**
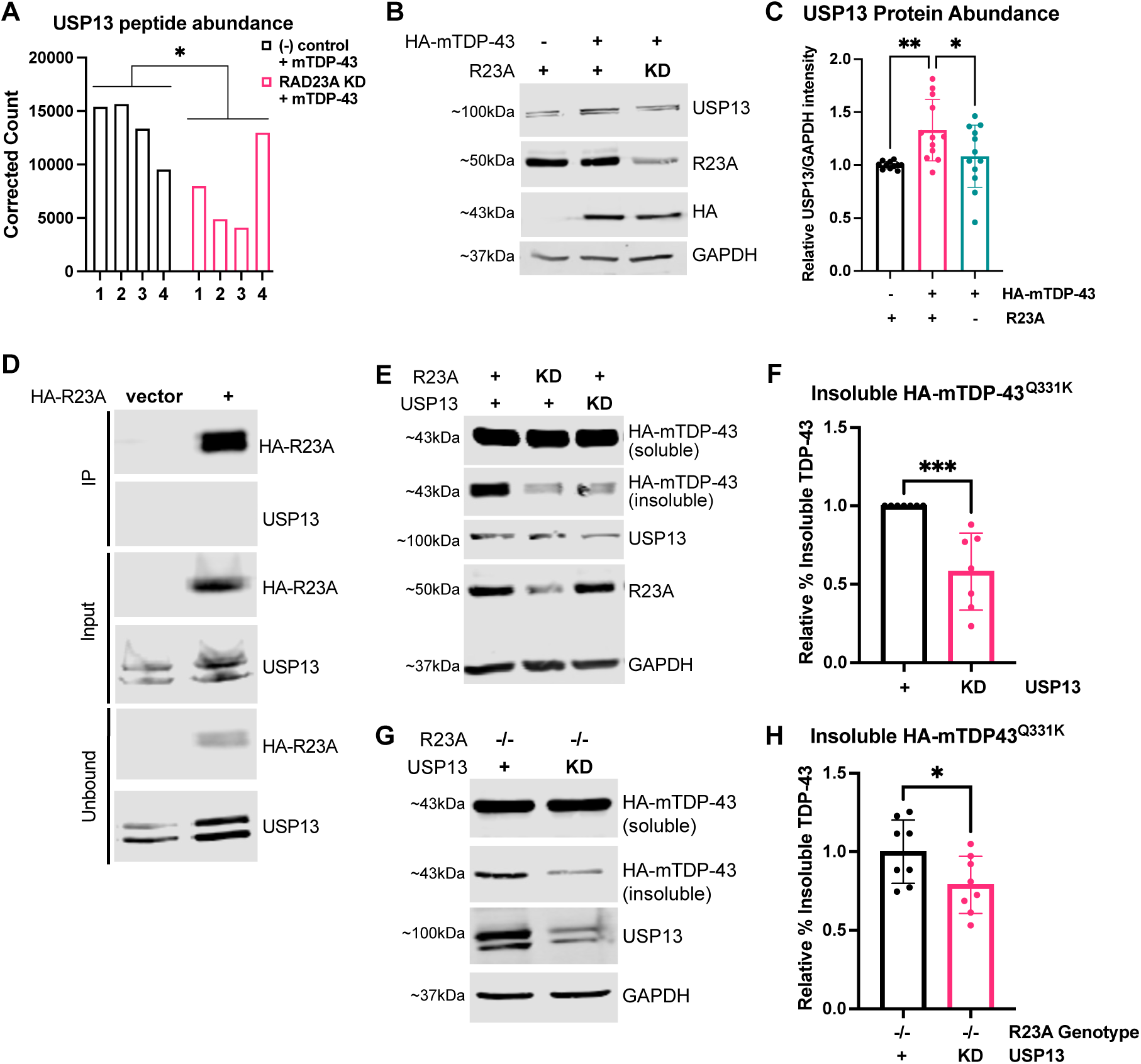
USP13 abundance is modulated by RAD23A and effects TDP-43 solubility potentially via a distinct pathway. **(A)** Corrected peptide counts for USP13 in +mTDP-43; R23A^CTRL^ and +mTDP-43; R23A^KD^, *p<0.05 of Log FC, one-tailed t-test. **B, C.** Immunoblot analysis of USP13 abundance following mTDP-43 induction or mTDP-43 induction with *RAD23A* knockdown. **(B)** Representative immunoblot of USP13 with indicated mTDP-43 status and siRNA treatment. **(C)** mTDP-43 induction increases USP13 abundance and *RAD23A* knockdown following mTDP-43 reduces USP13 abundance, one way ANOVA, F(2,32)=5.823, **p<0.01, represented relative to no mTDP-43 control siRNA. Data represent the mean +-SD, N=6 experimental replicates, 1-2 technical replicates. **(D)** Immunoblot of HA-RAD23A immunoprecipitation, showing no co-IP with USP13. **E, F.** Inducible HA-mTDP-43 cells were plated, transfected with indicated siRNA, induced with DOX for 24hrs, lysates collected and subjected to SarkoSpin fractionation. **(E)** *RAD23A* and *USP13* knockdown efficiency representative immunoblot and HA-mTDP-43 soluble and insoluble fractions following SarkoSpin fractionation representative immunoblot. **(F)** *USP13* knockdown reduces the percentage of HA-mTDP-43 present in the insoluble fraction (quantified as percent of total TDP-43) expressed relative to negative control. Data represent the mean +/−SD, N=7. (***p<0.001, two-tailed t-test. **G, H.** R23A^−/−^ HA-mTDP-43 inducible cells were plated, transfected with indicated siRNA, induced with DOX for 24hrs, collected and subjected to SarkoSpin fractionation. **(G)** *USP13* knockdown efficiency representative immunoblot and HA-mTDP-43 soluble and insoluble fractions following SarkoSpin fractionation representative immunoblot. **(H)** *USP13* knockdown further reduces the percentage of HA-mTDP-43 present in the insoluble fraction in RAD23A KO cells (quantified as percent of total TDP-43) expressed relative to negative control. Data represent the mean +/−SD, N=4, two technical replicates, *p<0.05, two-tailed t-test.

### USP13 knockdown in neurons reduces aggregates and improves survival

We then confirmed our TDP-43 solubility studies in primary rat cortical neuron cultures infected with human mTDP-43^A315T^ expressing virus, where *USP13* KD reduces the percentage of TDP-43 in the insoluble fraction by ∼46% (Fig. 4A and B). We next asked if the effects of *USP13* KD extended to aggregated wild-type TDP-43, as this is the prevalent species in patient autopsy tissue. In primary rat cortical neurons infected with wild type human TDP-43 expressing virus, we see a ∼67% reduction in the percentage of TDP-43 in the insoluble fraction (Fig. 4C and D).

**Figure 4.**
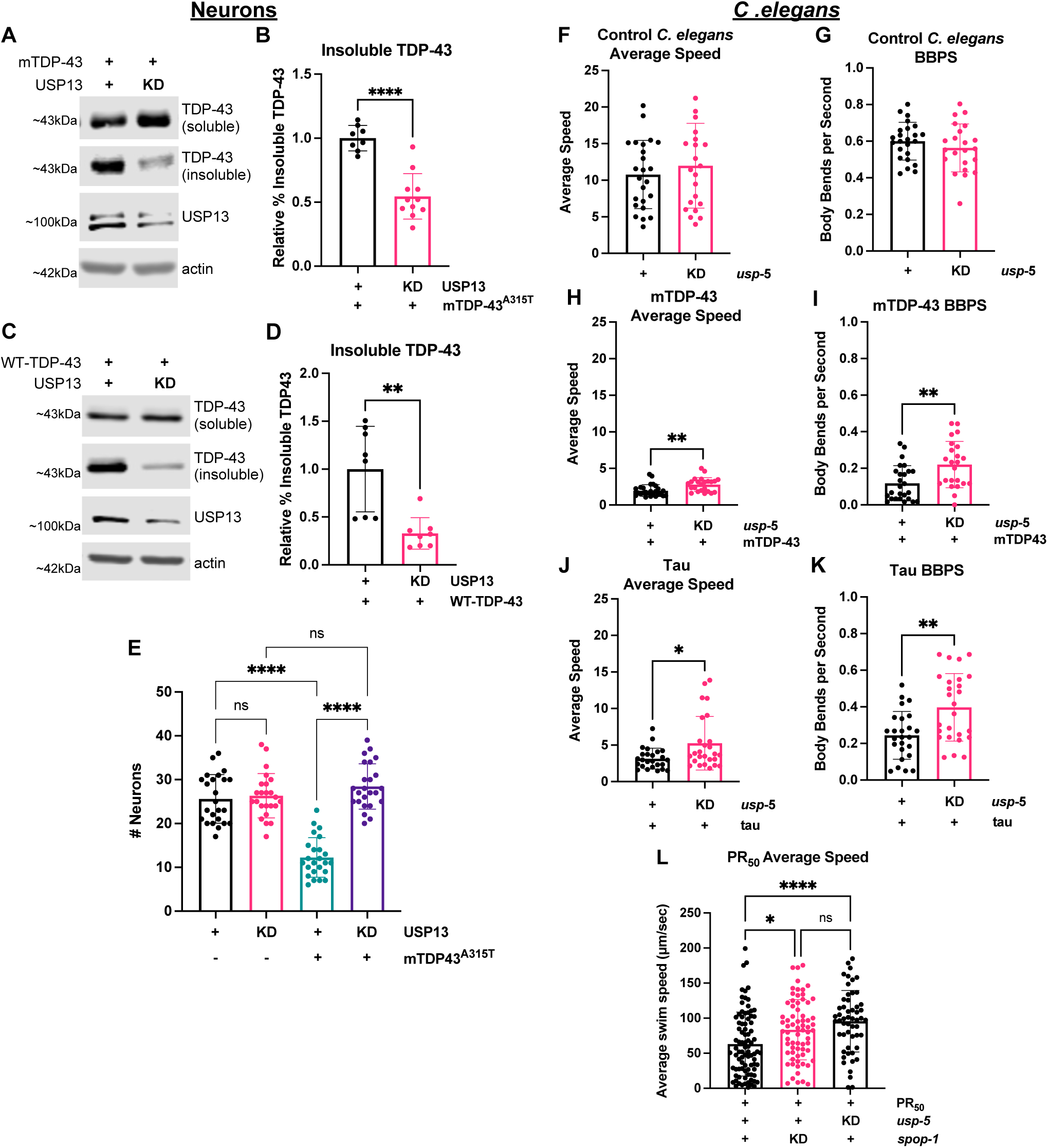
*USP13* knockdown reduces insoluble TDP-43, blunts TDP-43-induced neuronal toxicity, and improves locomotor deficits in multiple *C. elegans* models of neurodegenerative disease. **A-D**. Primary rat cortical neuron cultures were transfected with indicated siRNAs on DIV4, indicted disease protein was expressed on DIV12 and collected after 48 hrs for SarkoSpin fractionation. **A, B**: HSV-mTDP-43^A315T^ expression **(A)** *USP13* siRNA efficiency and TDP-43 soluble and insoluble fractions, representative immunoblot. **(B)** *USP13* knockdown reduces TDP-43 in the insoluble fraction (quantified as a percent of total TDP-43) relative to the negative control. Data represent the mean +/− SD N=4, 2-3 technical replicates (****p<0.0001, two-tailed t-test). **C, D.** HSV-WT-TDP-43 expression **(C)** *USP13* siRNA efficiency and TDP-43 soluble and insoluble fractions, representative immunoblot. **(D)** *USP13* knockdown reduces TDP-43 in the insoluble fraction (quantified as a percent of total TDP-43) relative to the negative control. Data represent the mean +/− SD N=4, 2 technical replicates (**p<0.01, two-tailed t-test). **(E)** USP13 blunts mTDP-43 induced neurotoxicity. Primary rat cortical neuronal cultures transfected with indicated siRNAs on DIV4, infected with HSV-mTDP-43^A315T^ or HSV-LacZ on DIV14, and fixed and stained with motor neuron marker SMI-32 on DIV19. N=4 experimental replicates, each data point is an area counted, N=6 areas per replicate. One-way ANOVA, F(5,138)=35.03,***p<0.0001. **F - L:** Quantification of locomotor function following feeding RNAi to the indicated gene in control or various disease models in *C.elegans.* **F, G.** Control RNAi sensitive, GRB73 animals, N=4, 5-8 animals per replicate. **(F)** Average speed, p=NS. **(G).** Body bends per second, BBPS, .p=NS. **H, I.** nervous system expression of mTDP43, CK674 N=4, 5-8 animals per replicate. **(H)** *usp-5* RNAi significantly increases average speed in mTDP-43 expressing animals, **p<0.01, two-tailed t-test. **(I)** *usp-5* feeding RNAi significantly increases BBPS in mTDP-43 expressing animals, **p<0.01,two-tailed t-test. **J, K**. Pan-neuronal over expression of human tau, CK144, N=4, 5-8 animals per replicate. **(J)** *usp-5* RNAi significantly increases average speed in human tau expressing animals,*p<0.05, two-tailed t-test. **(K)** *usp-5* feeding RNAi significantly increases BBPS in human tau expressing animals, **p<0.01,two-tailed t-test. **(L)** myo-3p:(PR)50::GFP expressing C.elegans, N=5, 50-100 animals per replicate. Both *usp-5(RNAi)* and *spop-1(RNAi)* significantly increase average swim speed in (PR)50 expressing animals p<0.0001, One-Way ANOVA with Tukey post hoc test

Next, we conducted a viability assay in primary rat motor neurons expressing human mTDP-43^A315T^ or a control via recombinant HSV vector. There are ∼50% less motor neurons remaining at DIV19 in cultures infected with A315T human mutant TDP-43 versus the control virus. KD of *USP13* causes a ∼132% increase in the number of motor neurons remaining at DIV19 in cultures infected with mTDP-43^A315T^ to numbers equal to that seen in cultures receiving control virus (Fig. 4E). *USP13* KD has no effect on the viability of neurons infected with the control virus.

### USP13 knockdown improves motor phenotypes in *C. elegans*

Encouraged by the *in vitro* studies, we asked whether reduction of *USP13* suppressed mutant TDP-43 induced phenotypes in an *in vivo* setting. *C.elegans* engineered to pan-neuronally express mutant TDP-43 display a severe uncoordinated (“Unc”) phenotype (32). We find that mutTDP-43 animals fed *usp-5(RNAi)* show a significant suppression of the locomotor deficit in both body bends per second (BBPS) (+87%) and average speed (+42%) (Fig. 4H and I). *usp*-5*(RNAi*) alone showed no significant differences in either BBPS or average speed (Fig. 4F and G). We then asked if reduction of *usp-*5 could suppress locomotor deficits evoked by a different pathological protein, tau. *USP13* KD has previously been shown to increase p-Tau ubiquitination in a mouse model that over-expresses human mutant amyloid precursor protein (33). *C. elegans* engineered to pan-neuronally over-express human tau display a profoundly uncoordinated phenotype. Feeding *usp-5(RNAi)* significantly suppresses tau-induced locomotor deficits in both BBPS (+63%) and average speed (+67%) (Fig. 4J and K). Finally, we asked if *usp*-5 KD could improve locomotion in a *C.elegans* model of the most common genetic cause of ALS, C9orf72 (34, 35). To do this we utilized animals with (PR)_50_ expression in the muscle cells with a completely penetrant developmental arrest and/or paralysis phenotype (32). Similar to the tdp-43 and tau models, *usp-5(RNAi)* suppressed the motility defects caused by (PR)_50._ We found a ∼52% increase in average swim speed, performing similarly to SPOP (36), a known modifier of DPR toxicity (Fig. 4L). In summary, we find that loss of *usp*-5 has broad beneficial effects on locomotor deficits on NDD models (e.g., tau, TDP-43, (PR)_50_) *in vivo*.

### USP13 complexes with but does not deubiquitinate TDP-43

Our results suggest USP13 modulates the toxicity of TDP-43 by influencing the aggregation dynamics that lead to the formation of insoluble TDP-43. As USP13 can function as a deubiquitinase, we hypothesized that under normal circumstances USP13 deubiquitinates TDP-43 and opposes its degradation. If this hypothesis is correct, we expect to see a biochemical interaction between USP13 and TDP-43 and KD of *USP13* would be expected to lead to increased ubiquitination of TDP-43. To determine if there is a physical interaction between USP13 and TDP-43, we conducted a co-IP using inducible HA-mTDP-43^Q331K^ or HA-WT-TDP-43 and immunoblotted for USP13. We find that USP13 does co-immunoprecipitate with both HA-mTDP-43^Q331K^ and HA-WT-TDP-43 (Fig. 5A and B). Additionally, we find that RAD23A does not co-immunoprecipitate with HA-mTDP-43^Q331K^ (Fig. 5C), indicating a further divergence in how the two proteins interact with and modulate TDP-43. This suggests that RAD23A may effect TDP-43 solubility indirectly or through a shared complex; whereas USP13 has a direct interaction with TDP-43 may effect TDP-43 solubility through direct stabilization of TDP-43.

**Figure 5.**
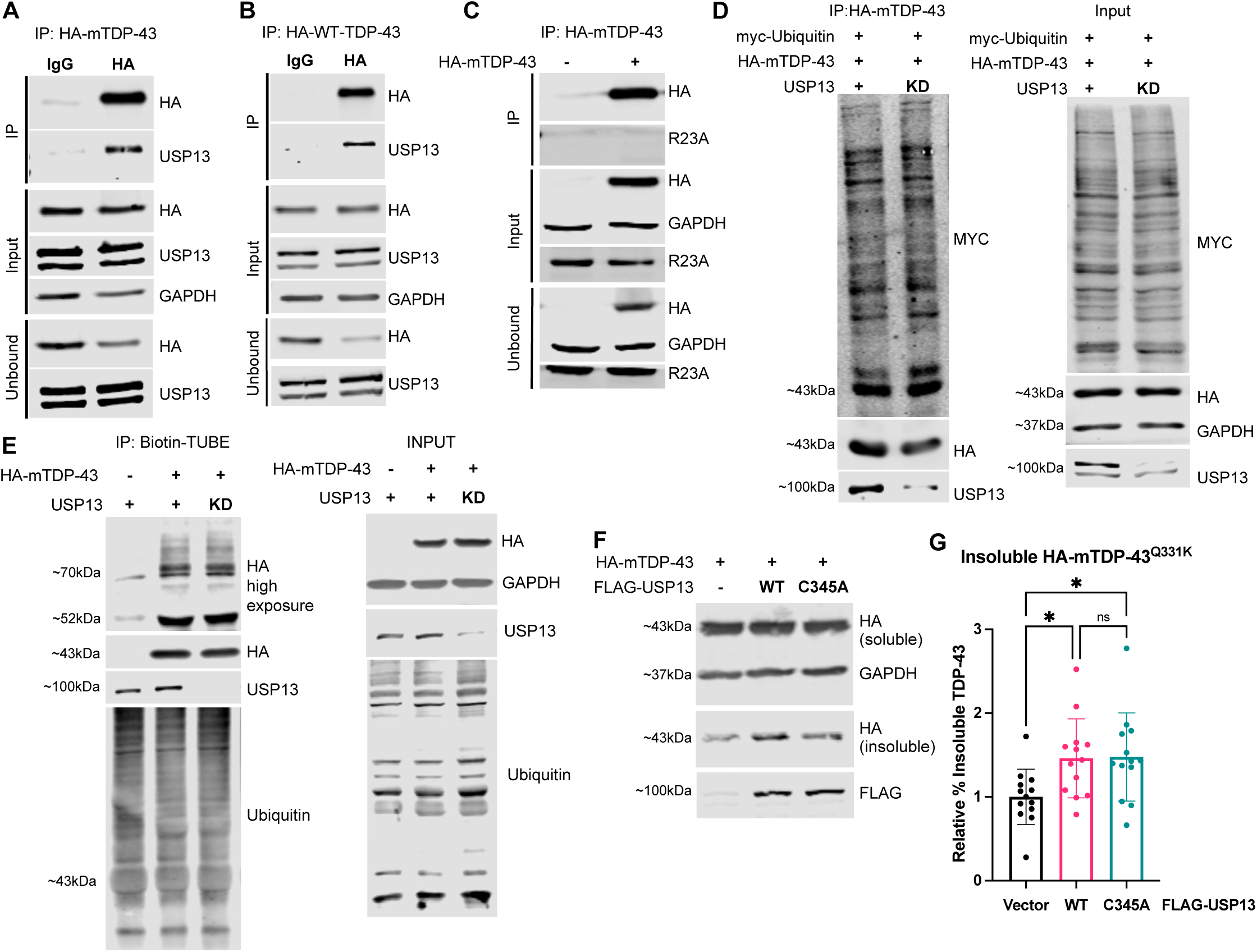
USP13 complexes with but does not deubiquitinate TDP-43 A-C. HA-mTDP-43 or HA-WT-TDP-43 HEK293 cells were induced for 24 hours, collected and lysed. Protein lysates were immunoprecipitated (IP) with anti-HA antibody and immunoblotted for indicated proteins. IP fraction (25% loaded), input (30 ug loaded), and unbound (∼5% loaded). **A.** Immunoblot following IP of HA-mTDP-43 showing co-IP of USP13. **B.** Immunoblot following IP of HA-WT-TDP-43 showing co-IP of USP13. **C.** Immunoblot following IP of HA-mTDP-43 showing no co-IP of RAD23A. **D.** HA-mTDP-43 HEK293 cells transfected to knockdown *USP13* and express myc-Ubiquitin, induced for 24 hours, collected and lysed. HA-mTDP-43 was immunoprecipitated. Immunoblot for myc-Ubiquitin shows no significant changes in abundance, indicating USP13 knockdown does not affect the ubiquitination of HA-mTDP-43. **E.** HA-mTDP-43 HEK293 cells transfected to knockdown *USP13*, induced for 24 hours, lysates were collected and Biotin-TUBE reagent was added. Biotin-TUBE was immunoprecipitated. Immunoblot for HA-mTDP-43 shows no significant changes in abundance, indicating *USP13* knockdown does not affect the ubiquitination of HA-mTDP-43. **F, G.** Inducible HA-mTDP-43^Q331K^ Flp-In T-REx HEK293 cells were plated, transfected with vector, FLAG-WT-USP13 or FLAG-USP13^C345A^ (catalytically inactive), induced with DOX for 24hrs, collected and subjected to SarkoSpin fractionation. **(F)** HA-mTDP-43^Q331K^ soluble and insoluble fractions and FLAG-USP13 abundance, representative immunoblot. **(G)** FLAG-WT-USP13 and FLAG-USP13^C345A^ significantly increases the percentage of HA-mTDP-43 in the insoluble fraction (quantified as a percent of total TDP-43) relative to negative control. Data represent the mean +/−SD, N=7, two technical replicates, one way ANOVA, F(2,36)=4.702, *p<0.05.

We then investigated the effect of USP13 on TDP-43 ubiquitination by *USP13* KD combined with exogenous expression of myc-Ubiquitin and induction of mTDP-43^Q331K^. When we immunoprecipitate HA-tagged TDP-43 and immunoblot for myc-Ubiquitin to assess the ubiquitination status of TDP-43, we find no significant changes in myc-Ubiquitin levels in either the input or IP fraction. This result indicates *USP13* KD does not alter ubiquitinated TDP-43 abundance (Fig. 5D). Additionally, we conducted the complementary experiment by application of Tandem Ubiquitin Binding Entity (TUBE) reagent, which can be used to enrich ubiquitinated proteins (37). Following *USP13* KD, induction of mTDP43, and IP of the Biotin-TUBE reagent, immunoblot for HA-mTDP-43 shows a clear ladder effect for poly-ubiquitinated TDP-43; however, we find that *USP13* KD does not change ubiquitinated HA-mTDP-43 levels or total ubiquitin levels in the input (Fig. 5E). Together these results are inconsistent with the hypothesis that USP13 is deubiquitinating TDP-43.

In order to investigate if the deubiquitinase activity of USP13 had an effect of TDP-43 aggregation, we exogenously expressed either empty vector, FLAG-USP13, or catalytically inactive FLAG-USP13^C345A^ and examined the effect on insoluble TDP-43 in our inducible model. We observe a significant increase in the percentage of insoluble TDP-43 when over-expressing wild-type FLAG-USP13 (46%) and catalytically inactive FLAG-USP13^C345A^ (47%) compared to empty vector (Fig. 5F and G). These results further suggest that USP13 influences the solubility of TDP-43 independent of its deubiquitinase activities.

## Discussion

Here we identify two modifiers of TDP-43 solubility and cytotoxicity. In both an inducible HEK cell line and primary neurons, we show that loss of RAD23A, but not RAD23B, leads to a decrease in TDP-43 accumulation into a detergent insoluble fraction and this is accompanied by a significant rewiring of the proteome. We demonstrate that reduced RAD23A abundance blunts the remodeling of the proteome induced by mTDP-43 expression. This is illustrated by the reduced abundance of USP13 following *RAD23A* KD in the presence of mTDP-43 induction. These findings proved to be of pathophysiological relevance as we find that *USP13* KD reduces insoluble TDP-43 in both an inducible model and primary neurons, blunts TDP-43-induced neurotoxicity in primary motor neurons, and reduces locomotor deficits in *C.elegans* models of NDDs. The mechanism for these improvements appears to be independent of the ubiquitination of total TDP-43. These results highlight the complex roles the ubiquitin proteasome system plays in maintaining proteostasis and overall cell health.

USP13, a deubiquitinase, plays a regulatory role in diverse cellular pathways such as autophagy (28, 38), the DNA damage response (29, 39), ER-associated degradation (ERAD) (40), energy metabolism(41), and innate immunity (42). USP13 has a complicated role in cancer prognosis and pathogenesis, with increased expression of USP13 reported to increase tumorigenesis and accelerate metastasis through stabilizing key substrates such as OGDH and ACLY, MCL-1, c-Myc in models of ovarian cancer (41), non-small cell lung cancer (43), cervical cancer (44), and glioblastoma (45). However, the tumor suppressor PTEN is also a direct substrate of USP13 and through these effects USP13 can also work to inhibit tumorigenesis in certain cancers such as breast carcinoma and bladder cancer (46, 47). These studies reveal the context specific actions of USP13 in maintaining cellular homeostasis.

More recently, a role for USP13 in neurodegeneration has been reported. USP13 regulates alpha-synuclein and p-tau ubiquitination and *USP13* KD increases alpha-synuclein and tau clearance in models of Parkinson’s and Alzheimer’s disease (33, 48). This enhanced clearance of aggregation prone proteins following *USP13* KD is in line with our findings for TDP-43. Parkin ubiquitination was also demonstrated to be regulated by USP13, with KD of *USP13* increasing parkin ubiquitination. Ubiquitination is known to be critical for activation of parkin’s E3 ligase activity, (49) and removal of this ubiquitination may have deleterious effects on downstream targets beyond alpha-synuclein. USP13 binding has been shown to stabilize another E3 ligase, Siah2, through binding via its ubiquitin associated (UBA) domains without deubiquitinating the substrate (50). This finding highlights that USP13 can function to stabilize a protein and interfere with its functions, through mechanisms independent of its canonical deubiquitinase activity.

While USP13 regulates the degradation of many aggregation prone proteins implicated in neurodegenerative disease, the effect of USP13 on TDP-43 appears to be distinct. We hypothesize that USP13 stabilizes TDP-43 without deubiquitination, potentially by binding the protein and preventing other protein-protein interactions that could lead to degradation of TDP-43. This hypothesis would be aligned with USP13’s actions on Siah2. In support of this, we show that TDP-43 and USP13 do physically interact, an observation consistent with a prior two-hybrid interaction screen in yeast that identified USP13 as a potential interactor of mutant TDP-43 (51); however, future work is needed to determine which domain of USP13 facilitates this interaction and if this interaction is critical for our observed reductions in insoluble TDP-43. In conclusion, we present USP13 as a new potent modifier of TDP-43 toxicity with its KD leading to reduced insoluble TDP-43, enhanced neuronal viability, and improved locomotor phenotypes. Therefore, targeting USP13 may have promising therapeutic potential in TDP-43 proteinopathies.

## Materials and Methods

### Antibodies

The following antibodies were used in this study: RAD23A (Abcam; ab108592), RAD23A (Cell Signaling; 24555S), RAD23B (Abcam; ab86781), HA (Cell Signaling; C29F4), GAPDH (Sigma; G8795), Actin (Novus; NB600-501), USP13 (Proteintech; 16840-1-AP), TDP-43 (Proteintech; 10782-2-AP), ubiquitin (Enzo; BML-PW0930-0100), Myc (Cell Signaling; 2278S), FLAG (Cell Signaling; 2368S), PBXIP1 (Proteintech; 12102-1-AP), CHCHD10 (Proteintech; 25671-1-AP), NDUFS5 (Proteintech; 15224-1-AP)

### Silencer Select siRNAs

The following siRNAs were used in this study: Negative control #1 (human, rat; 4390843), RAD23A (human; s11729), RAD23A (rat; s167251); RAD23B (human; s11732); RAD23B (rat; s220827), USP13 (human; s117130); USP13 (rat; s160289),PBXIP1 (human; s32944), CHCHD10 (human; s226550), NCEH1 (human; s33292); NDUFS5 (human; s9397), TPP1 (human; s502514).

### HA-mTDP-43^Q331K^ Flp-In T-Rex cell line creation

#### Plasmid generation

A gBlock insert containing 15 bp overhangs for the EcoRV site of pcDNA5/FRT/TO (Invitrogen™ V652020), an N-terminal HA-tag and the full cDNA of human TDP-43 with a Q331K point mutation was purchased as an IDT gBlock. This insert was this cloned into the EcoRV site of pcDNA5/FRT/TO using In-Fusion Snap Assembly master mix according to the manufacturer’s protocol (Takara 638951).

#### Cell culture, stable cell line creation

HEK293 Flp-In T-REx cells (ThermoFisher R78007) were cultured in DMEM (Gibco™ 11965092), 10% Tetracycline free FBS (Gibco™ A4736201), 1x penicillin-streptomycin (Gibco™ 15140122), 1X GlutaMAX™ (Gibco™ 35050061), 15 ug/mL Blasticidin S (Gibco™ A1113903) and 100 ug/mL zeocin (Gibco™ R25001). To create a stable cell line expressing HA-mTDP-43^Q331K^, 3×10^5^ cells were plated per well of a 6-well plate and transfected when 70% confluent 48 hours later with 125 ng of HA-mTDP-43-pcDNA5/FRT/TO, 1.125 ug pOG44 (Invitrogen™ V600520), and 6.25 uL Lipofectamine 2000 (Invitrogen™ 11668500) per well. After 48 hours, cells were re-plated in media without zeocin and with 150 ug/mL hygromycin B (Gibco™ 10687010) for selection. Cells were selected for 3 weeks before re-plating for expansion and screening for successful integration via immunoblotting.

**HA-WT-TDP-43 Flp-In T-REx cells** are from the lab of Dr. Magdalini Polymenidou (13).

### R23A^+/−^ and R23A^−/−^ CRISPR cell line creation. crRNA selection, knockout cell line creation

*Rad23a* knockout cell lines were generated in the Flp-In T-REx HEK cell line. Pre-designed crRNA guides to RAD23A (Hs.Cas9.RAD23A.AC and Hs.Cas9.RAD23A.AD), tracrRNA (1073189), and Cas9 nuclease (1081058) were purchased from IDT. Selected crRNA guides were duplexed with tracrRNA and combined Cas9 nuclease before transfection with RNAiMAX lipofectamine. Single colonies were isolated, expanded, genomic DNA was isolated via Direct PCR lysis (Viagen; 301-C), and sequenced for successful edits.

### siRNA KD and doxycycline induction in Flp-In T-REx HEK293 cells

Briefly, doxycycline-inducible Flp-In T-REx HEK293 cells were plated on rat tail collagen, transfected at ∼60-70% confluence with RNAiMAX and silencer select siRNAs to KD the gene of interest, and the next day HA-mTDP-43 expression was induced at a final concentration of 5 ng/mL of doxycycline (DOX) for 24 hours.

### SarkoSpin Fractionation

Sarkospin fractionation was carried out as described in (13). 2 wells of a 6 well plate of HEK293 or primary neuronal cultures were collected by scraping in ice cold PBS, pelleted by centrifugation, and resuspended in lysis buffer containing 0.5% sarkosyl and benzonase. Protein concentration was measured and equalized before proceeding with sarkosyl solubilization and centrifugation to isolate insoluble material.

### Rat Primary Neuron Culture

Mixed spinal cord neuron cultures were prepared as described previously described (52). Briefly, an astrocyte feeder layer was prepared from the cortex of newborn Sprague Dawley rat pups (postnatal day 2, P2) and grown to ∼80% confluency. Subsequently, dissociated embryonic day 15 (E15) spinal cord neurons were added. One to two days later, AraC (5 mm) (catalog #C6645; Sigma) was added for 24 h to arrest astrocyte proliferation. Cultures were maintained in glia-conditioned medium supplemented with the following trophic factors (1.0 ng/ml each): human neurotrophin-3, human neurotrophin-4, human brain-derived neurotrophic factor, and rat ciliary neurotrophic factor (Alomone Labs). Half of the culture medium was replaced on a biweekly basis.

### siRNA KD and HSV infection in rat neuronal cultures

Primary rat cortical neuronal cultures were transfected on DIV 4-5 using RNAiMAX lipofectamine and Silencer Select siRNAs in to KD the gene of interest. In order to over-express human TDP-43 or mutant TDP-43^A315T^, primary rat cortical neurons were infected with herpes simplex viruses (NeuroTools) at 1-2 uL/mL expressing the human protein at DIV 11-14. Cells were collected for fractionation and/or immunoblot 48 hours later.

### Rat Motor Neuron Survival Assay

Primary rat spinal cord neurons were plated on a feeder layer of astrocytes. On DIV 4, neurons were transfected with siRNA as described above and maintained until DIV 14. On DIV 14 neurons were infected with mTDP-43^A315T^-HSV at 0.75 uL/mL and maintained until DIV19. On DIV19 cultures were fixed, stained with SMI-32 motor neuron marker and neurons were counted.

### HA-mTDP-43^Q331K^ Immunoprecipitation

HA-mTDP-43^Q331K^ HEK293 cells were plated in a 10 cm dish, induced with DOX at 5 ng/uL for 24 hours prior to lysis in 250 uL Pierce Lysis Buffer, protease inhibitor cocktail, and N-ethylmaleimide. IP was conducted using Pierce Anti-HA Magnetic Beads (Thermo Scientific #88836), mouse IgG1 monoclonal antibody HA-epitope YPYDVPDYA. Pierce Anti-c-Myc magnetic mouse IgG1 monoclonal antibody beads were used as the IgG control. 1 mg of protein was loaded onto the beads for each preparation, IP was conducted according to manufacturer’s protocol with 25% of the eluted IP fraction loaded for immunoblot. Co-IP for each protein assayed was determined via immunoblot. For De-ubiquitination measurements cells were also transfected with 8 μg myc-ubiquitin and lipofectamine 2000 prior to DOX induction.

### TMT-based quantitative proteomics

#### Sample prep for MS Analysis

HA-mTDP-43^Q331K^ HEK293 cells were plated in a 10 cm dish, transfected with negative control no.1 siRNA (ThermoFisher) or *RAD23A* siRNA using RNAiMAX lipofectamine, followed by induction with DOX at 5 ng/uL for 24 hours if indicated. Cells were then washed PBS, and dry 10 cm dishes were stored in −80C until all replicates were complete. After all replicates were complete, samples were lysed on ice in 300 uL RIPA buffer with protease inhibitors (Sigma, P8340). Sample concentration was determined by BCA (Fisher Scientific) and 1 mg aliquots were prepared. Protein was precipitated using methanol chloroform precipitation. Protein pellets were resuspended in 8 M urea (Thermo Fisher Scientific, 29700) prepared in 100 mM ammonium bicarbonate solution (Fluka, 09830). Dithiothreitol (DTT, DOT Scientific Inc, DSD11000) was applied to a final concentration of 5 mM. After incubation at RT for 20 mins, iodoacetamide (IAA, Sigma-Aldrich, I1149) was added to a final concentration of 15 mM and incubated for 20 mins at RT in the dark. Excess IAA was quenched with DTT for 15 mins. Samples were diluted with 100 mM ammonium bicarbonate solution, and digested for three hrs with Lys-C protease (1:100, Thermo Fisher Scientific, 90307_3668048707) at 37°C. Trypsin (1:100, Promega, V5280) was then added for overnight incubation at 37°C with intensive agitation (1000 rpm). The next day, reaction was quenched by adding 1% trifluoroacetic acid (TFA, Fisher Scientific, O4902-100). The SEC fraction samples were desalted using ZipTip (Thermo Fisher–Pierce, 87784). All samples were vacuum centrifuged to dry.

#### Tandem Mass Tag Labeling

Our protocol was based on previously reported methods (53). C18 column-desalted peptides were resuspended with 100 mM HEPES pH 8.5 and concentrations were measured by micro BCA kit (Fisher Scientific, Cat# PI23235). For each sample, 100 μg of peptide labeled with TMT reagent (0.4 mg, dissolved in 40 μl anhydrous acetonitrile, Thermo Fisher Scientific, A44520) made at a final concentration of 30% (v/v) acetonitrile (ACN). Following incubation at RT for 2 hrs with agitation, hydroxylamine (to a final concentration of 0.3% (v/v)) was added to quench the reaction for 15 min. TMT-tagged samples were mixed at a 1:1:1:1:1:1:1:1:1:1:1:1:1:1:1:1 ratio. Combined sample was vacuum centrifuged to dry, resuspended, and subjected to HyperSep C18 Cartridges (ThermoFisher, 60108-302).

#### Peptide Fractionation

We used a high pH reverse-phase peptide fractionation kit (Thermo Fisher Scientific, 84868) to get eight fractions (5.0%, 10.0%, 12.5%, 15.0%, 17.5%, 20.0%, 22.5%, 25.0% and 50% of ACN in 0.1% triethylamine solution). The high pH peptide fractions were directly loaded into the autosampler for MS analysis without further desalting.

#### Tandem Mass spectrometry

3 μg per fraction or sample were auto-sampler loaded with an UltiMate 3000 HPLC pump onto a vented Acclaim Pepmap 100, 75 um x 2 cm, nanoViper trap column coupled to a nanoViper analytical column (Thermo Fisher, 164570, 3 µm, 100 Å, C18, 0.075 mm, 500 mm) with stainless steel emitter tip assembled on the Nanospray Flex Ion Source with a spray voltage of 2000 V. An Orbitrap Fusion (Thermo Fisher) was used to acquire all the MS spectral data. Buffer A contained 94.785% H_2_O with 5% ACN and 0.125% FA, and buffer B contained 99.875% ACN with 0.125% FA. For TMT-MS experiments, the chromatographic run was for 4 hrs in total with the following profile: 0-7% for 7, 10% for 6, 25% for 160, 33% for 40, 50% for 7, 95% for 5 and again 95% for 15 mins respectively. For other MS experiments, the chromatographic run was for 2 hours in total with the following profile: 2–8% for 6, 8–24% for 64, 24–36% for 20, 36–55% for 10, 55–95% for 10, 95% for 10 mins. We used a multiNotch MS^3^-based TMT method to analyze all the TMT samples (53–55). The scan sequence began with an MS^1^ spectrum (Orbitrap analysis, resolution 120,000, 400-1400 Th, AGC target 2×10^5^, maximum injection time 200 ms). MS^2^ analysis, ‘Top speed’ (2s), Collision-induced dissociation (CID, quadrupole ion trap analysis, AGC 4×10^3^, NCE 35, maximum injection time 150 ms). MS^3^ analysis, top ten precursors, fragmented by HCD prior to Orbitrap analysis (NCE 55, max AGC 5×10^4^, maximum injection time 250 ms, isolation specificity 0.5 Th, resolution 60,000). We used CID-MS^2^ method for other experiments as previously described. Briefly, ion transfer tube temp = 300 °C, Easy-IC internal mass calibration, default charge state = 2 and cycle time = 3 s. Detector type set to Orbitrap, with 60K resolution, with wide quad isolation, mass range = normal, scan range = 300-1500 m/z, max injection time = 50 ms, AGC target = 200,000, microscans = 1, S-lens RF level = 60, without source fragmentation, and datatype = positive and centroid. MIPS was set as on, included charge states = 2-6 (reject unassigned). Dynamic exclusion enabled with n = 1 for 30 s and 45 s exclusion duration at 10 ppm for high and low. Precursor selection decision = most intense, top 20, isolation window = 1.6, scan range = auto normal, first mass = 110, collision energy 30%, CID, Detector type = ion trap, OT resolution = 30K, IT scan rate = rapid, max injection time = 75 ms, AGC target = 10,000, Q = 0.25, inject ions for all available parallelizable time.

#### MS data analysis and quantification

Protein ID/quantification and analysis were performed with Integrated Proteomics Pipeline-IP2 (Bruker, Madison, WI. http://www.integratedproteomics.com) using ProLuCID (56, 57), DTASelect2 (58, 59), Census and Quantitative Analysis (For TMT-MS experiments). Spectrum raw files were extracted into MS1, MS2 and MS3 (For TMT experiments) files using RawConverter (http://fields.scripps.edu/downloads.php). The tandem mass spectra were searched against UniProt human (downloaded on 5-18-2023) protein databases (60) and matched to sequences using the ProLuCID/SEQUEST algorithm (ProLuCID version 3.1) with 5 ppm peptide mass tolerance for precursor ions and 600 ppm for fragment ions. The search space included all fully and half-tryptic peptide candidates within the mass tolerance window with no-miscleavage constraint, assembled, and filtered with DTASelect2 through IP2. To estimate peptide probabilities and false-discovery rates (FDR) accurately, we used a target/decoy database containing the reversed sequences of all the proteins appended to the target database (60). Each protein identified was required to have a minimum of one peptide of minimal length of six amino acid residues; however, this peptide had to be an excellent match with an FDR < 1% (< 5% for TiO_2_ column-enriched samples) and at least one excellent peptide match. After peptide/spectrum matches were filtered, we estimated that the peptide FDRs were ≤ 1% for each sample analysis. Resulting protein lists include subset proteins to allow for consideration of all possible protein forms implicated by at least two given peptides identified from the complex protein mixtures. We used Census and Quantitative Analysis in IP2 for protein quantification of TMT-MS experiments and protein quantification was determined by summing all TMT report ion counts. For TMT experiments: static modification: 57.02146 C for carbamidomethylation, 304.2071 for 16-plex TMT tagging; differential modifications: 304.2071 for N-terminal 16-plex TMT tagging, 42.0106 for N-terminal Acetylation. Resulting protein lists include subset proteins to allow for consideration of all protein isoforms implicated by at least two given peptides identified from protein mixtures (61).

#### TMT Final Data analysis

Proteins were removed from analysis that did not have at least two successful replicates in each group within a given comparison. P-value was calculated using a 1-tailed t-test. Log_2_FC was calculated based on the mean of each comparison group. Pathway analysis was conducted using Metascape gene analysis tools (21).

### USP13 Plasmid Generation

FLAG-USP13 and FLAG-USP13^C345A^: gBlock inserts containing 15 bp overhangs for the EcoRV site of p1006 (gift from Dr. Rachel Neve, Harvard Medical School) and N-terminal FLAG-tag and the full cDNA of human USP13 or full cDNA of human USP13 with the C345A mutation were purchased as IDT gBlocks. These inserts were cloned into the EcoRV site of p1006 using In-Fusion Snap Assembly master mix according to the manufacturer’s protocol (Takara 638951).

### Deubiquitination Assay by Tandem Ubiquitin Binding Entity (TUBEs) and Streptavidin IP

HA-mTDP-43^Q331K^ HEK293 cells were plated, transfected, DOX induced and lysed as above for HA-IP. TUBE reagent was added to IP lysis buffer (LifeSensors; UM101:TUBE1). IP was conducted using Dynabeads^TM^ MyOne^TM^ Streptavidin C1 (ThermoFisher; 65001). 1 mg of protein was loaded for each prep, IP was conducted according to manufacturer’s protocol with 25% of the eluted IP fraction for immunoblot. Ubiquitination was measured by subsequent immunoblot for HA-mTDP-43 in the eluted IP fraction.

### C. elegans strains

All animals were grown on nematode growth media (NGM) plates seeded with *Escherichia coli* OP50 and maintained at either 20°C or 15°C. CK674, CK144, and GRB73 were obtained from the laboratory of Dr. Brian C. Kraemer and maintained at 15°C. PR_50_ animals (strain OG755 *drIs34 [myo-3p::(PR)50-GFP; myo-3p::dsRed2]*) are from the laboratory of Todd Lamitina.

### RNAi Feeding and Locomotor Assay

*C.elegans* strain CK674 (expressing neuronal TDP-43), CK144 (pan-neuronal over-expression of human tau), and GRB73 (RDE-1 background RNAi sensitive control) were used for RNAi feeding and locomotor assays. RNAi bacteria was prepared from the Ahringer library Plate #8, E6. (T27A3.2)) (*usp*-5). 4-6 pregnant animals were placed onto IPTG plates with RNAi bacteria, allowed to lay eggs, and those eggs were allowed to develop to L4 stage over 4-5 days at 15C. 5-8 L4 animals were then moved onto an agar plate with no OP50 into a drop of M9 buffer (22 mM KH2PO4, 42mM Na_2_HPO_4_, 86 mM NaCl). Swimming was recorded for 30 seconds using a Zeiss Stemi SV11 stereomicroscope and Lumenera Infinity 2 camera and videos were processed using the wrMTrck plugin (62) in ImageJ to calculate average speed and body bends per second (BBPS).

### PR_50_ *C.elegans* Locomotor Assay

For the DPR toxicity assays, strain OG755 *drIs34 [myo-3p::(PR)50-GFP; myo-3p::dsRed2]* was utilized. Eight gravid adults grown on *gfp(RNAi)* plates were transferred to either *empty vector(RNAi)* or *usp-5(RNAi)* (Clone T27A3.2 from MRC Geneservice library) plates, as previously described (63). After 48 hours, L4 stage animals were picked to new RNAi plates and placed at 25°C. 72 hours later, Day 3 adults were moved into a customized recording chamber (64) with 80µl of M9 buffer and gently covered with a glass coverslip. 30s of swimming behavior was captured at 14 frames per second and animals analyzed using the Wormlab system (MBF Bioscience). Only animals with a track time of ≥20 seconds were used for analysis. Average swim speed (µm/sec) was analyzed, and outlier data were removed using the ROUT function in GraphPad Prism (Q=1%). The average of the resulting data (n=50-100 animals per genotype) was analyzed using one-way ANOVA and Tukey’s multiple comparison test (for 3 or more groups) or an unpaired T-test (for 2 groups). P-values of <0.05 were considered significant.

### Statistical Analysis

We used Microsoft Office Excel and GraphPad Prism version 10.1.1 to analyze data and plot graphs. Data were expressed as mean ± S.D unless specifically noted as S.E.M. For comparisons among 2 or more groups, ordinary one-way ANOVA was used; and for comparisons between 2 groups, two-tailed, unpaired Student *t*-tests were used, unless specifically noted. P≤0.05 was considered significant.

## Acknowledgments

We would like to express sincere appreciation to Dr. Daniel J. Finley for his thoughtful review of this work. This work was supported by the Assistant Secretary of Defense for Health Affairs endorsed by the Department of Defense, through the Amyotrophic Lateral Sclerosis Research Program Therapeutic Idea Award under Award No. (HT94252411080). Opinions, interpretations, conclusions and recommendations are those of the author and are not necessarily endorsed by the Assistant Secretary of Defense for Health Affairs endorsed by the Department of Defense. C.D., J.M.P, and R.G.K were supported by the US Public Health Service (NS05225 and NS087077), the Heather Koster Family Charitable Fund, and the Les Turner ALS Foundation; J.N.S., Y.-Z.W. and K.K.G were supported by the US Public Health Service (S10OD032464); N.S. was supported by the US Public Health Service (NS122257); T.L. and C.S. were supported by the US Public Health Service (NS124802).

## Author Contributions

C.D., T.L., J.N.S., and R.G.K designed research; C.D., J.M.P, N.S., C.S., K.K.G., and Y.Z.W. performed research; C.D., N.S., C.S., K.K.G., and Y.Z.W. analyzed data; C.D. and R.G.K wrote the paper.

