## Supplemental Figures 1 to 5 for "Ubiquitin Proteasome System Components, RAD23A and USP13, Modulate TDP-43 Solubility and Neuronal Toxicity"

**Figure S1: *RAD23A* and *RAD23B* knockdown reduce the amount of cytosolic mTDP-43.**

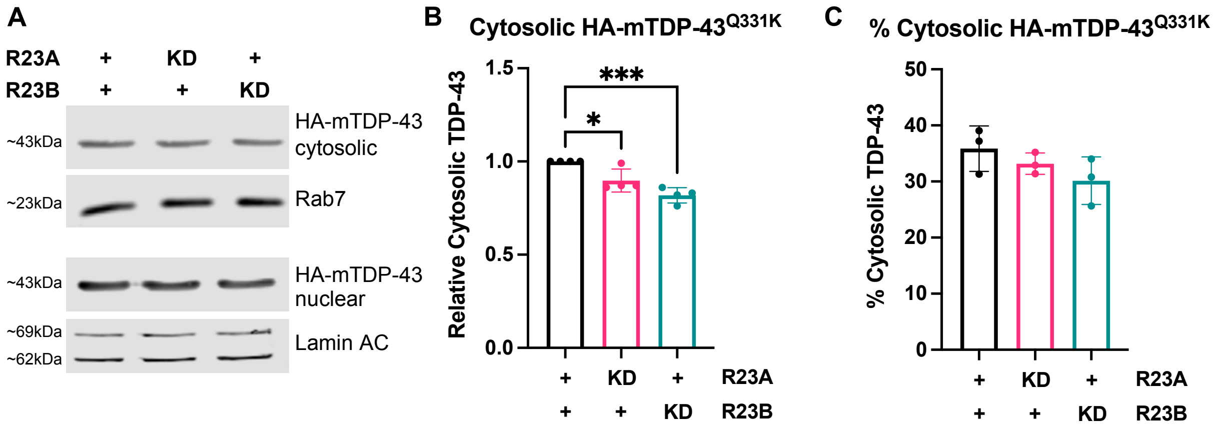


**Figure S1. *RAD23A* and *RAD23B* knockdown reduce the amount of cytosolic mTDP-43.** Inducible HA-mTDP-43 Flp-In T-REx HEK293 cells were plated, transfected with indicated siRNA, induced with DOX for 24hrs, collected and subjected to nuclear cytosolic fractionation. **(A)** HA-mTDP-43 cytosolic and nuclear fractions, representative immunoblot. **(B)** *RAD23A* and *RAD23B* knockdown reduce the amount of HA-mTDP-43 present in the cytosolic fraction relative to negative control, normalized to Rab7 loading control. Data represent the mean +/-SD, N=4, one way ANOVA, $F(2,9)=18.12$, ***$p<0.001$ **(C)** Neither *RAD23A* or *RAD23B* knockdown reduces the percentage of cytosolic HA-mTDP-43 relative to the nega.tive control. Cytosolic normalized to Rab7, nuclear normalized to Lamin A/C. Data represent the mean +- SD, N=3, one way ANOVA, $F(2,6)=1.938$, p=NS.

#### Figure S2. Venn diagrams showing the shared proteins between the each comparison group with direction of change in expression included

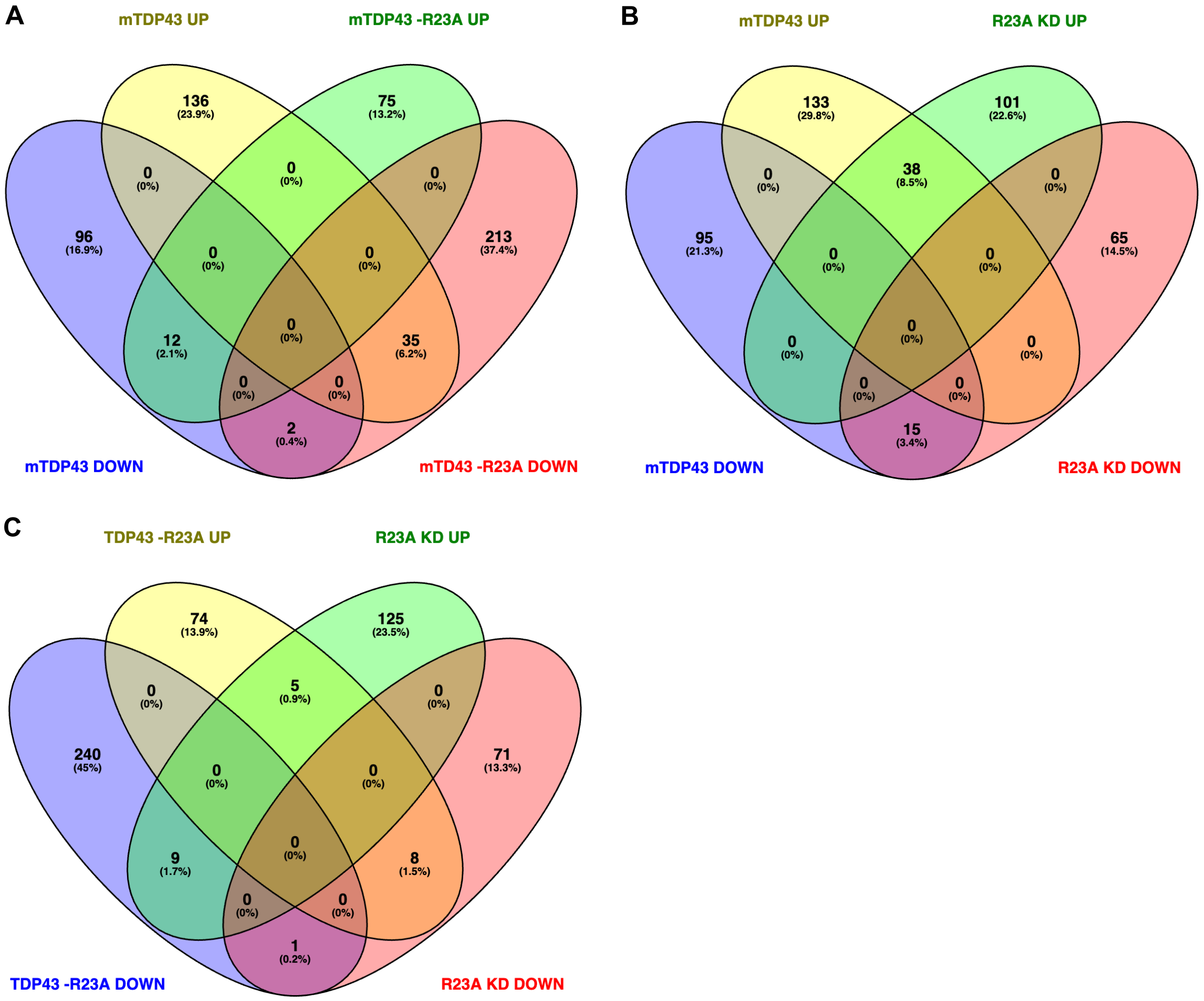


**Figure S2. Venn diagrams showing the shared proteins between the each comparison group with direction of change in expression included.** **(A)** Group 1, proteins altered by mTDP-43 induction (upregulated and downregulated) versus Group 3, proteins altered by *RAD23A* knockdown in the presence of mTDP-43 (upregulated and downregulated). **(B)** Group 1, proteins altered by mTDP-43 induction (upregulated and downregulated) versus Group 2, proteins altered by *RAD23A* knockdown (upregulated and downregulated). **(C)** Group 2, proteins altered by *RAD23A* knockdown (upregulated and downregulated) versus Group 3, proteins altered by *RAD23A* knockdown in the presence of mTDP-43 (upregulated and downregulated).

### Figure S3: Pathways enriched in Group 1 and Group 2

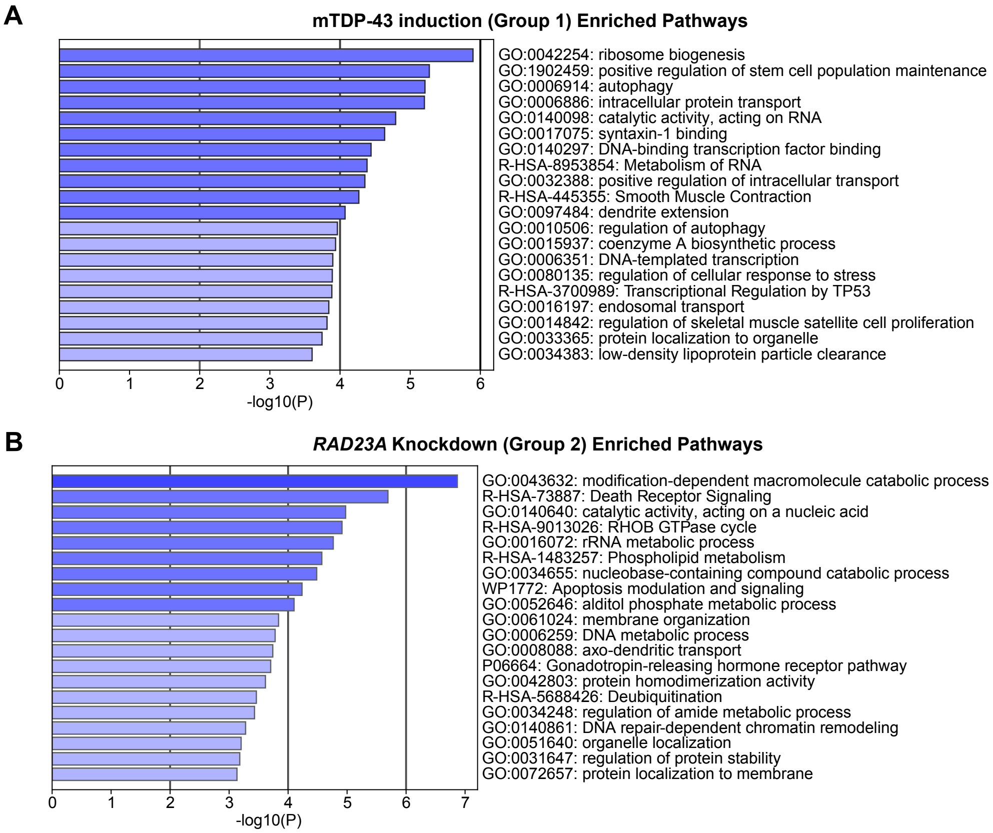


**Figure S3. Pathways enriched in Group 1 and Group 2. A - B.** Pathway analysis conducted on proteins with a $p<0.05$ and Log2FC>0.25 or $Log_2FC<-0.25$. **(A)** Pathways enriched in Group 1 (by mTDP-43 induction) using Metascape gene analysis, proteins with both increased and decreased expression included in analysis. **(B)** Pathways enriched in Group 2 (by *RAD23A* knockdown) using Metascape gene analysis, proteins with both increased and decreased expression included in analysis.

#### Figure S4: Analyis of shared pathways between the three analysis groups.

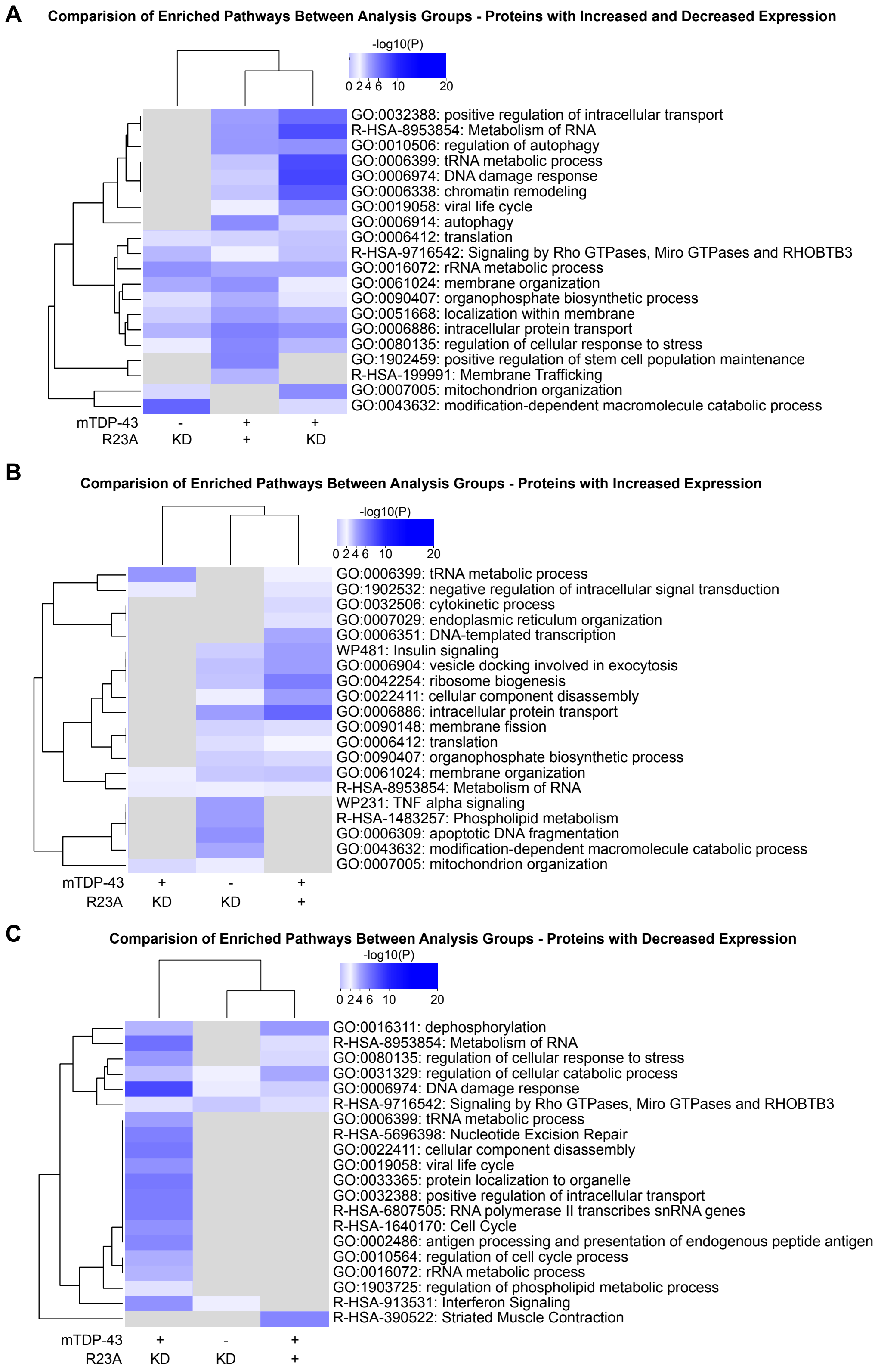


**Figure S4. Comparison of shared pathways between the three analysis groups. A - C:** Pathway analysis conducted on proteins with a p<0.05 and $Log_2FC>0.25$ or $Log_2FC<-0.25$. **(A)** Group 1, Group 2, and Group 3 comparison of shared pathways with proteins with both increased and decreased expression included in analysis. **(B)** Group 1, Group 2, and Group 3 comparison of shared pathways with proteins of increased expression included in analysis. **(C)** Group 1, Group 2, and Group 3 comparison of shared pathways with proteins of decreased expression included in analysis.

#### Figure S5: Screening of Candidates Identified from Proteomics

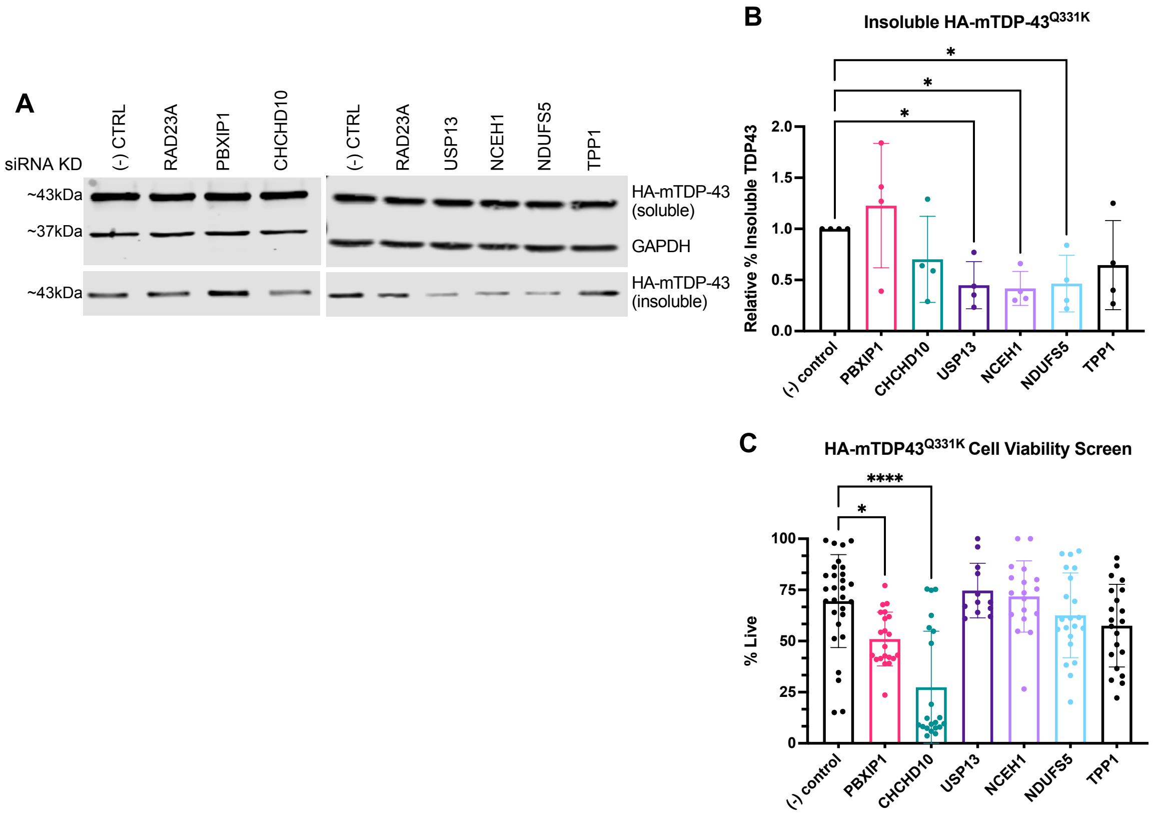


**Figure S5. Screening of Candidates Identified from Proteomics A - B:** Inducible HA-mTDP-43 Flp-In T-REx HEK293 cells were plated, transfected with indicated siRNA, induced with DOX for 24hrs, collected and subjected to SarkoSpin fractionation. **(A)** HA-mTDP-43 soluble and insoluble fractions following knockdown of indicated siRNA, representative immunoblot. **(B)** *USP13*, *NCEH1*, and *NDUFS5* knockdown reduce the percentage of HA-mTDP-43 present in the insoluble fraction (quantified as a percent of total TDP-43) relative to negative control. Data represent the mean +/-SEM, N=4, one way ANOVA, $F(6,21)=2.971$, *$p<0.05$ **(C)** Percent of cells staining positive for calcein after saturated DOX induction for 72 hours and indicated siRNA knockdown, Data represent the mean +/-SEM, N= 2-4, 6-8 technical replicates per experiment, one-way ANOVA, $F(8,184)=9.645$, ****$p<0.0001$

#### Supplementary Data Table 1
see attached excel file

**Table S1:** TMT-based quantitivate proteomics full protein list with individual replicate counts, fold change and p-value; Group 3 proteins with individual replicate counts, fold change and p-value; Group 2 proteins with individual replicate counts, fold change and p-value; Group 2 proteins with individual replicate counts, fold change and p-value;

#### Supplementary Data Table 2
see attached excel file

**Table S2:** Metascape pathway analysis pathway identifiers, enrichment, and protein lists used for Fig 2F; Fig 2G; Fig 2H; Fig S3A; Fig S3B; Fig S4A; Fig S4B; Fig S4C
